# Federated Learning for Predicting Compound Mechanism of Action Based on Image-data from Cell Painting

**DOI:** 10.1101/2024.02.09.579629

**Authors:** Li Ju, Andreas Hellander, Ola Spjuth

## Abstract

Having access to sufficient data is essential in order to train accurate machine learning models, but much data is not publicly available. In drug discovery this is particularly evident, as much data is withheld at pharmaceutical companies for various reasons. Federated Learning (FL) aims at training a joint model between multiple parties but without disclosing data between the parties. In this work, we leverage Federated Learning to predict compound Mechanism of Action (MoA) using fluorescence image data from cell painting. Our study evaluates the effectiveness and efficiency of FL, comparing to non-collaborative and data-sharing collaborative learning in diverse scenarios. Specifically, we investigate the impact of data heterogeneity across participants on MoA prediction, an essential concern in real-life applications of FL, and demonstrate the benefits for all involved parties. This work highlights the potential of federated learning in multi-institutional collaborative machine learning for drug discovery and assessment of chemicals, offering a promising avenue to overcome data-sharing constraints.

## 1 Introduction

Artificial Intelligence (AI) and Machine Learning (ML) are seeing increased adoption in the field of drug discovery and toxicology [1], providing researchers with the capability to predict e.g. chemical and biological properties [2], pharmacokinetics [3] as well as toxicity and safety [4]. By integrating AI and ML into the drug discovery processes, the potential exists to significantly reduce both the time and speed of developing novel therapeutics [1]. Another related area AI holds high potential is chemical risk assessment [5]. Recently, Deep Learning in the form of Deep Neural Networks (DNN) has advanced the capabilities of applications relying on various types of imaging [6, 7]. The opportunities provided by cloud computing has also contributed significantly to this development by providing rapid access to large-scale computational resources and software frameworks for modeling [8]. These technologies pave the way for enhanced precision and effectiveness, initiating a new era of possibilities in pharmaceutical research and development and assessment of chemicals.

Having access to sufficient data is essential in order to train accurate machine learning models. However, the majority of the world’s data is not readily available to the data scientist. In drug discovery this is particularly evident, as much data is withheld at pharmaceutical companies, for reasons including data sensitivity (e.g. connected to patients and clinical trials), intellectual property protection, regulatory compliance, the potential value of novel compounds in future applications, and to gain commercial advantages towards competitors or an unwillingness to let competitors gain insights into ongoing and past projects [9, 10, 11].

While drug discovery is a highly competitive industry, many problems are shared among organizations, such as predicting chemical properties, toxicity and the biological mechanisms of drug leads. As a consequence, many experiments are repeated among companies and academic laboratories. Companies and researchers are increasingly recognizing the value of data sharing and collaboration in accelerating drug discovery and development, and several initiatives have been established to promote and advance data sharing, including IMI OpenPHACTS [12], and PistoiaAlliance (https://www.pistoiaalliance.org). Despite this trend the main bulk of a pharmaceutical company’s databases are not shared with other parties.

Federated Machine Learning, or simply Federate Learning (FL), is a decentralized machine learning paradigm that enables collaborative model training across a network of participants. Privacy is preserved by keeping data of participants local [13]. This technique empowers a central server to refine a cohesive model through the progressive aggregation of locally-trained models. While federated learning has found applications across domains such as Natural Language Processing [14], Computer Vision [15], and Internet of Things [16, 17], its potential extension into broader fields remains an open question. Significant challenges confront the practical application of federated learning, notably the inherent heterogeneity in both data quality and distribution [18]. Such discrepancies can potentially undermine the overall performance of the model.

In recent years there have been reported several success stories of federated learning in medical applications, such as for distinguishing healthy brain tissue from tissue affected by cancer cells from MRI images [19] and for diagnosing leukemia from blood transcriptomes and tuberculosis and lung pathologies from X-ray images [20], and predicting clinical outcome in COVID-19 patients from electronic medical records and X-ray images [21]. These implementations relies to a large extent on deep neural networks in the form of convolutional neural networks. In drug discovery, FL has been demonstrated primarily for predictions on chemical structure [22, 23, 6]. The wide-spread data type (chemical structure, represented by atoms and bonds e.g. in SMILES format) and the simplicity of numerical representation (chemical descriptors) makes the problem readily accessible for federated learning. A remaining challenge is the heterogeneity in experimental assays to determine ground truth for properties in the training sets, and how to integrate data from different assays or when transitioning between assays [24].

In drug discovery, profiling cells using high-content imaging is increasingly used to capture the morphological effect upon exposure to compounds [25, 26]. To this end, Deep Learning methods have shown great results [27, 28, 29]. The most common method for multiplexed cell profiling is Cell Painting [30] where cells are stained with mutliple dyes (normally 6) targeting different organelles, and imaged in multiple (normally 5) channels. Several deep learning methods have been used to analyze cell painting data in applications including predicting lung cancer variants [31], for chemical hazard evaluation [32], studying combination effects of compounds [33], diagnosing Parkinson’s disease [34] and for antiviral research [35]. However, federated learning applied to image-based cell profiling in a drug discovery context has not been reported.

In drug discovery, predicting the mechanism of action (MoA) of drugs and new drug leads constitute an important but challenging task [36]. MoA refers to the specific biochemical interaction through which a drug substance produces its pharmacological effect. In essence, it’s the process by which a molecule, such as a drug, induces a change in a biological target, typically a protein, that leads to a therapeutic outcome. The majority of efforts to predict MoA relies on ground truth from specific in vitro assays for individual MoA classes, with features from chemical structure [36, 37], gene expression [38], proteomics [39], and cell morphology [40, 41]. Understanding the mechanism of action is key in drug development as it, apart from explaining the drugs primary interaction, also helps predict potential side effects, drug-drug interactions, and provides a rational basis for therapeutic use. In this manuscript we apply federated learning to the problem of predicting drug MoA based on high-content imaging of cells exposed to drug perturbations according to the Cell Painting protocol. Specifically, we assess the effectiveness and efficiency of federated learning against non-collaborative learning and data-sharing collaborative learning in various scenarios. We particularly study the impacts of data heterogeneity for MoA prediction, a major concern for federated learning in real-life applications, and empirically demonstrate the benefits of federated learning.

## 2 Methods

### 2.1 Data Description

We utilized a dataset of cell painting images of fluorescence microscopy originating from a previous study [42] with data deposited at Figshare [43]. It consists of 231 drugs, each annotated with one out of 10 mechanism-of-action labels (see Table 1). Briefly, the human osteosarcoma cell line U2OS was exposed to drugs and incubated for 48 hours in microplates having 384 wells each. Each compound was tested in six replicate wells, stained using 6 dyes according to the Cell Painting protocol [30], and images of five channels were captured using wide-field microscopy with a resolution of 2160 *×* 2160 from nine different sites per well. The wells were distributed across 18 different plates, designed using PLAID (Plate Layouts using Artificial Intelligence Design [44]). In total, the dataset consists of 13878 images. For more detailed information on experimental protocols, see [42].

**Table 1:** The dataset consists of 231 drugs, each labeled with a single mechanism-of-action (MoA) describing how the drug interacts with cells. ‘Num. Images’ corresponds to the number of independent images captured from wet-lab experiments where U2OS cells were exposed to each drug and stained and imaged according to the Cell Painting protocol [30].

| MoA | Encoding | Num. Compounds | Num. Images |
| --- | --- | --- | --- |
| retinoid receptor agonist | 0 | 19 | 1026 |
| protein synthesis inhibitor | 1 | 23 | 1242 |
| topoisomerase inhibitor | 2 | 32 | 1728 |
| Aurora kinase inhibitor | 3 | 20 | 1080 |
| ATPase inhibitor | 4 | 18 | 1026 |
| HSP inhibitor | 5 | 24 | 1296 |
| HDAC inhibitor | 6 | 33 | 1782 |
| JAK inhibitor | 7 | 21 | 1188 |
| PARP inhibitor | 8 | 21 | 1134 |
| tubulin polymerization inhibitor | 9 | 20 | 1080 |

### 2.2 Problem Formulation & Optimization Method

The aim of the study is to train a joint model to predict the MoA of a compound, based on cell painting images where the U2OS cell line has been treated with this compound. In our setting, we construct different federated learning scenarios with multiple parties, each of which only has access to its private set of cell painting data. The objective is to compare the performance of a federated model compared to models trained on each party’s private data independently, and to the model trained with all data pooled (i.e. centralized data).

MoA prediction from distributed fluorescence image data can be formulated as a federated optimization problem. Assume that there exists an unknown conditional probability mass function *p*(*y*|***x***) where ***x*** is the fluorescence image and *y* is the MoA of compounds. We want to approximate *p*(*y*|***x***) with a neural network *f* (***x***; ***θ***) parameterized by ***θ***, with *K* sets of data 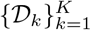 where 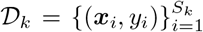 With a predefined loss function *ℓ*(*ŷ, y*), we have the optimization problem:

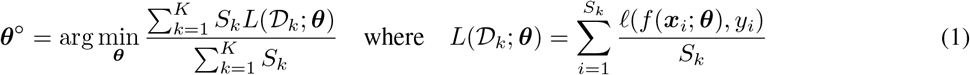

`FedAvg` is the de facto algorithm to solve the federated optimization problem [13]. The pseudo-code for FedAvg is shown in Algorithm 1.

#### Algorithm 1

`FedAvg`

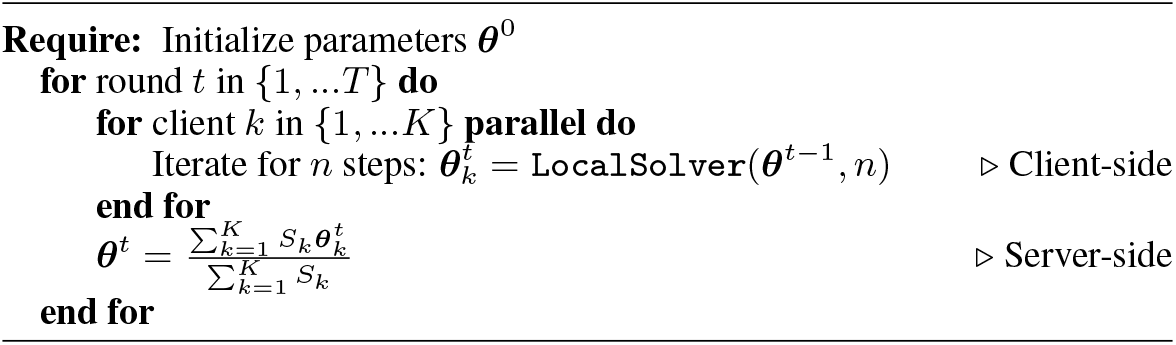

### 2.3 Experiment Setup

In drug discovery and for chemical safety assessment, it is common that companies and organizations have performed similar experiments, such as biological assays to measure safety endpoints, but on different chemical compounds. At the same time, most organizations incorporate publicly available data, e.g. from ChEMBL [45] into their internal databases to enrich their datasets. We here construct three hypothetical scenarios where organizations collaborate to train a shared model to predict 10 different labels (MoAs) but with different distributions of the available data:

- Uniform: Participants have access to private data of similar sizes covering all MoAs.
- Unbalanced: Similar to Uniform but participants have different data sizes for the MoAs.
- Non-IID: Similar to Uniform but only one client (Specialized) has access to private data on one or more specific MoAs.

The Uniform and Unbalanced scenario attempts to resemble a collaboration with common endpoints, such as produced with well-established assays based on OECD guidelines, and where the data content is either distributed uniformly or unbalanced between the participants. The Non-IID scenario resembles acollaboration where one partner has performed specific assays for a set of compounds, such as an advanced or novel mechanistic or safety test.

To set up these scenarios, we generate distributed datasets as illustrated in Figure 1. The collected data were divided into a total of *K* + 2 partitions, with one serving as a public/shared partition, another as the test partition, and the remaining partitions as private datasets. Each clients local data is then composed of its private data partition augmented with the partition simulating the public/shared dataset. Importantly, each client retains access exclusively to their respective local datasets.

**Figure 1.**
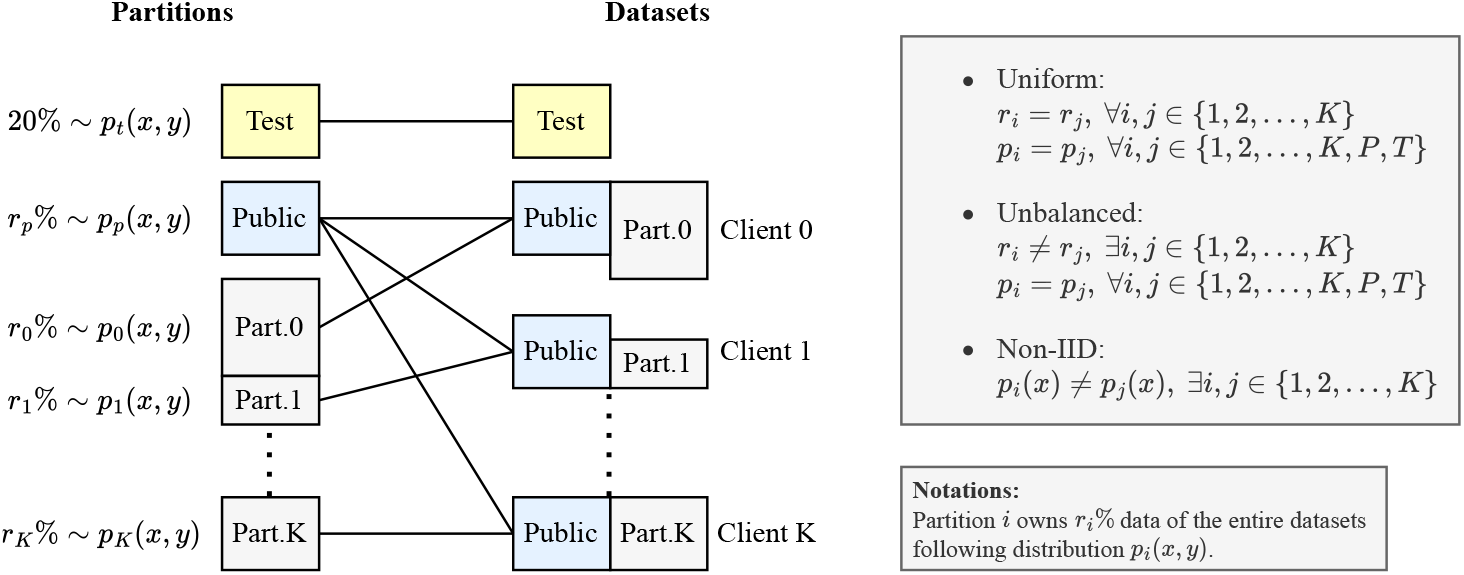
Partitioning schemes for generating distributed datasets for the three scenarious (uniform, unbalanced and non-IID). Each setup begins by dividing the available data into a total of *K* + 2 partitions, comprising one test partition, one public/shared partition, and *K* private partitions. The *K* + 2 partitions are combined to create one test dataset and *K* local datasets that combine individual private partitions with the public shared partition. **Uniform:** In the uniform scenario, data of 20% and *r*_*p*_% randomly-sampled compounds serve as the test and public partitions, respectively. The remaining compounds are uniformly distributed among *K* private partitions. **Unbalanced:** In the unbalanced scenario, mirroring the uniform setup, data of 20% and *r*_*p*_% randomly-sampled compounds are designated as the test and public partitions. The remaining data is distributed among *K* partitions with varying sample sizes per partition but following identical distributions. **Non-IID:** In the non-IID scenario, 20% of the data constitutes the test partition, with deliberate skews in the MoA distributions of data for other partitions, to ensure that data distributions between client datasets are inherently different.

The evaluation of trained models is carried out using the test set. Notably, the partitioning is conducted at the compound level, ensuring that samples of a given compound are not distributed across different clients.

### 2.4 Machine Learning

#### 2.4.1 Model Architecture

The goal of this study is not to achieve the highest possible prediction accuracy through exploration of various neural network models. Rather, our aim is to demonstrate the effectiveness and improvement of federated learning over centralized learning. To this end, we utilize AlexNet [46] and VGG13 [47] as practical choices to illustrate the effectiveness of federated learning for two different model architectures. We have adjusted AlexNet and VGG13 to match the depth and resolution of images in our dataset. The model architectures are shown in Figure 2. Throughout we use cross-entropy as loss function.

**Figure 2.**
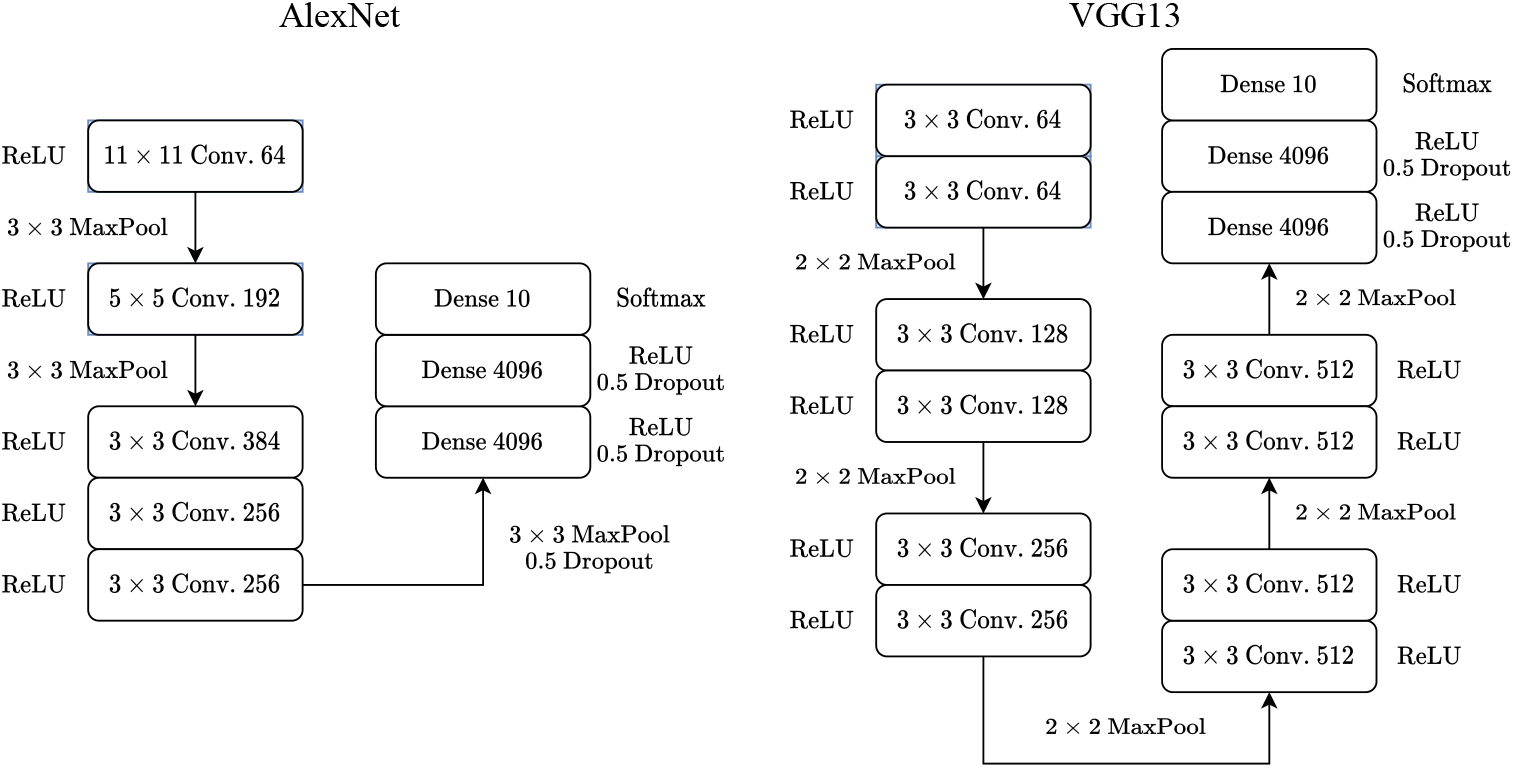
Architectural Layout of the base models we use for federared learning experiments. Both models employ a sequence of convolutional layers to encode features from fluorescence images into a latent space. Subsequently, a sequence of dense layers is employed for classifying these encodings. **Left:** AlexNet **Right:** VGG13

#### 2.4.2 Data Preprocessing and Augmentation

The image data are first normalize to reduce plate-level noise by subtracting the mean pixel intensities of control DMSO wells in each plate. Subsequently, all five-channel images are resized to 256 *×* 256 pixels from their original resolution and standardized to have zero mean and unit variance as 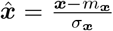 where *m*_***x***_ and *σ*_***x***_ are channel-wise mean and standard deviation of ***x*** in local datasets respectively.

To enhance the robustness and generalization ability of the model, we employed data augmentation techniques [48]. The images were randomly flipped and/or rotated by 90^*?*^ before being fed to the neural network for training. For test data, no data augmentation is done for preprocessing, only data normalization is employed.

#### 2.4.3 Comparison to Centralized and Local Training

We compare two baseline training settings to federated training:

- Centralized Learning (CL): Here all local datasets are pooled together to train a model using centralized learning. This is seen as the best possible model that can be obtained from existing distributed datasets if clients would give their permission to share data with a central actor.
- Local Learning (LL): Here each participant trains their own model using only their local dataset. These models are the best models an individual client can archive using their private datasets. Based on these we can evaluate the improvement potential by federated training.

We fine-tuned the hyperparameters for centralized learning and local learning using grid search. For all tasks, we used momentum SGD optimizer with a learning rate of *η* = 0.01 and momentum factor *m* = 0.9. Models were trained for 100 epochs with a fixed batch size of 30 for centralized learning and 10 for local learning. Exponential learning rate decay was applied with a decay rate of *γ* = 0.96 to improve model convergence. Gradient clipping [49] was used with a maximum *l*_2_ norm of gradients of value 3.0 to prevent gradient explosion.

#### 2.4.2 Federated Training

The federated models are trained using FEDn [50], a production-grade federated learning framework deployed on NAISS cloud [51], UPPMAX [52] and Alvis [53] High-Performance Computing (HPC) clusters in a distributed manner. The architecture of the training framework is shown in Figure 3. The models are trained using the FedAvg algorithm, as shown in Algorithm 1.

**Figure 3.**
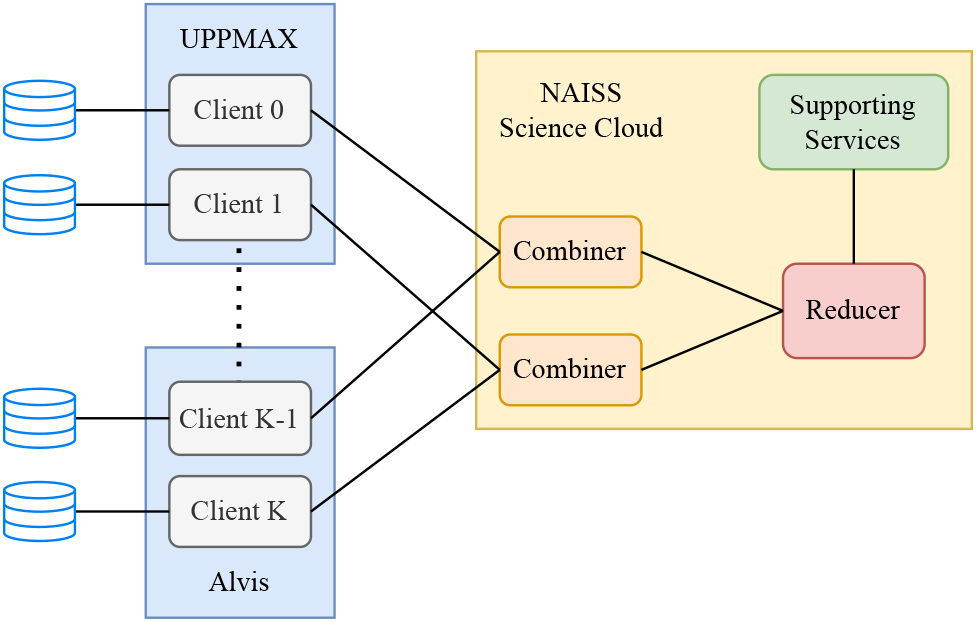
Training Framework Architecture. The training framework, FEDn, is geographically distributed across three High-Performance Computing (HPC) clusters: NAISS Science Cloud, UPPMAX, and Alvis, located in Umeå, Uppsala, and Gothenburg, Sweden, respectively. Within this framework, components in the NAISS Science Cloud are responsible for aggregating local models in a hierarchical manner, supported by essential services. Local training occurs on HPC clusters equipped with GPU resources, specifically UPPMAX Snowy and Alvis. The partitioned data are stored on the respective HPC clusters and can only be accessed by their assigned owners.

The hyperparameters for used for training are shared across all clients and are fine-tuned to obtain the best performance. For all setups, the local optimizer is momentum SGD, with learning rate *η* = 0.01 and momentum factor *m* = 0.9. Decay of local learning rate is applied to improve the convergence of the federated training with *γ* = 0.96, as suggested in [13]. In total, federated models were trained for 100 global rounds, with local models being trained for 1 epochs per round with a batch size of 20.

All experiments, including local training, centralized training and federated training, are repeated three times from models randomly initialized with different random seeds. Means and standard deviations of metrics are reported.

## 3 Experimental Results

### 3.1 FL Outperforms Local Learning and is Comparable to Centralized Learning

#### 3.1.1 Average Accuracy

We first assess the predictive performance of models trained under different schemes, measuring prediction accuracy on the test dataset in the uniform and unbalanced scenarios. To compare federated models and local and centralized models quantitatively, we define the metric Relative Change 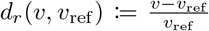. Student’s *t*-test is applied to determine whether the prediction accuracy of federated models is significantly higher than that of local models and lower than that of centralized models. The comparison of prediction accuracy of different models and their relative changes are shown in Figure 4. The results for the *t*-test is reported in Table 2.

**Table 2:** Results of Student’s *t*-Test: Comparing Model Performance. Student’s *t*-test is employed to assess whether the federated models outperform their corresponding local models and underperform centralized models with statistical significance.

| $H_0$ | Uniform | | Unbalanced | |
| --- | --- | --- | --- | --- |
|  | AlexNet | VGG13 | AlexNet | VGG13 |
| $\text{Acc}_{\text{LL},0} < \text{Acc}_{\text{FL}}$ | 0.005 | 0.000 | 0.005 | 0.005 |
| $\text{Acc}_{\text{LL},1} < \text{Acc}_{\text{FL}}$ | 0.004 | 0.002 | 0.000 | 0.000 |
| $\text{Acc}_{\text{LL},2} < \text{Acc}_{\text{FL}}$ | 0.002 | 0.003 | 0.001 | 0.000 |
| $\text{Acc}_{\text{CL}} > \text{Acc}_{\text{FL}}$ | 0.091 | 0.264 | 0.611 | 0.531 |

**Figure 4.**
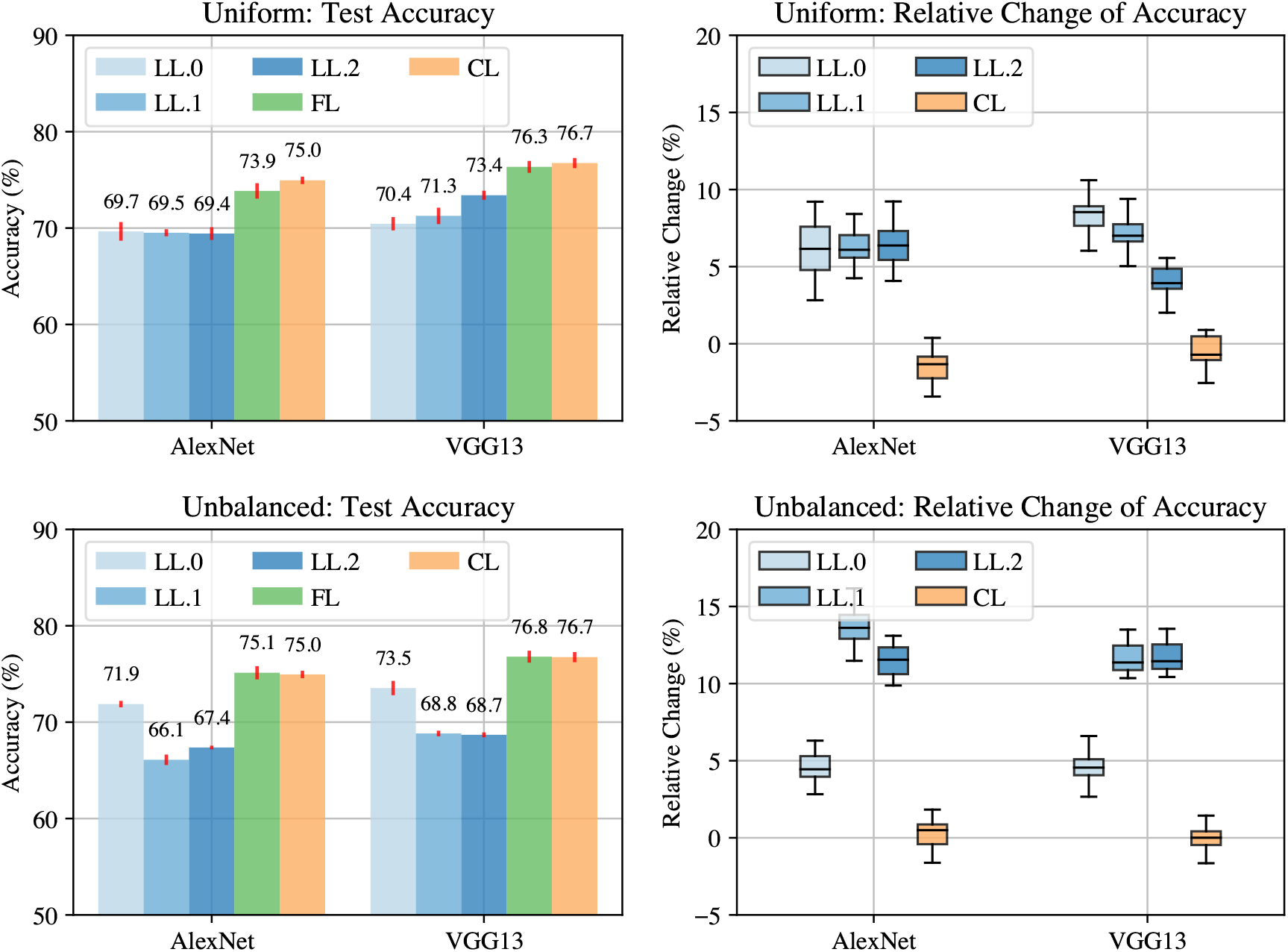
Comparative Analysis of Model Performance. This figure compares experimental results of models trained under different schemes in both uniform and unbalanced setups. Local learning on client *m*, federated learning and centralized learning are denoted as LL.*m*, FL and CL, respectively. **Left**: Prediction accuracy of AlexNet and VGG13 models on the test dataset, trained using different learning schemes for both uniform and unbalanced setups. **Right**: Relative change in prediction accuracy on the test dataset for federated models compared to their corresponding local and centralized models for both uniform and unbalanced setups.

In the uniform setup, federated and centralized AlexNet models achieve a mean prediction accuracy with standard deviation of 73.9 *±*0.3% and 75.0 *±*0.1%, respectively on the test dataset, while the prediction accuracy of the best-performing local model falls below 69.7%. In terms of relative changes in prediction accuracy, the federated AlexNet models exhibit an approximately 7% improvement over their local counterparts. Results from a *t*-Test indicate that AlexNet models trained using federated learning significantly outperform those trained with local learning. Also,federated AlexNet models do not significantly underperform centralized AlexNet models in terms of prediction accuracy. Similar results are observed when VGG13 is served as the base model. The results for the uniform setup are visualized in the upper section of Figure 4.

In the unbalanced setup, where Client 0 has the most data while Client 1 and 2 have less, this data distribution is reflected in the prediction accuracy of their corresponding local models. Despite the varying sizes of local datasets, the federated model consistently has significantly better prediction accuracy than local models. Additionally, it is observed that the less private data a client possesses, the more improvement could be gained from federated learning, with relative changes ranging from 4.6% to 13.4% on average. It is also observed that federated models and centralized models have similar predicting performance for both AlexNet and VGG13 models. The results for the unbalanced setup are visualized in the lower section of Figure 4.

Overall empirical results consistently show that, in either the uniform or unbalanced setup and when using either the AlexNet or VGG13 model, federated learning outperforms local learning and exhibit prediction accuracy comparable to centralized learning. Additionally, it’s noteworthy that the standard deviations of test accuracy for federated models are similar to those of local models, indicating that federated training is able to train models with stable performances from distributed data.

#### 3.1.2 Precision and Recall for Individual MoAs

The improvement in average accuracy doesn’t necessarily guarantee an improvement in the prediction performance of each individual MoAs. This is referred to as the problem of fairness in machine learning models [54, 55]. We here compare precision and recalls of models for individual MoAs. Figure 5 compares AlexNet models trained with federated, local and centralized learning for each specific MoA in both uniform and unbalanced setups.

**Figure 5.**
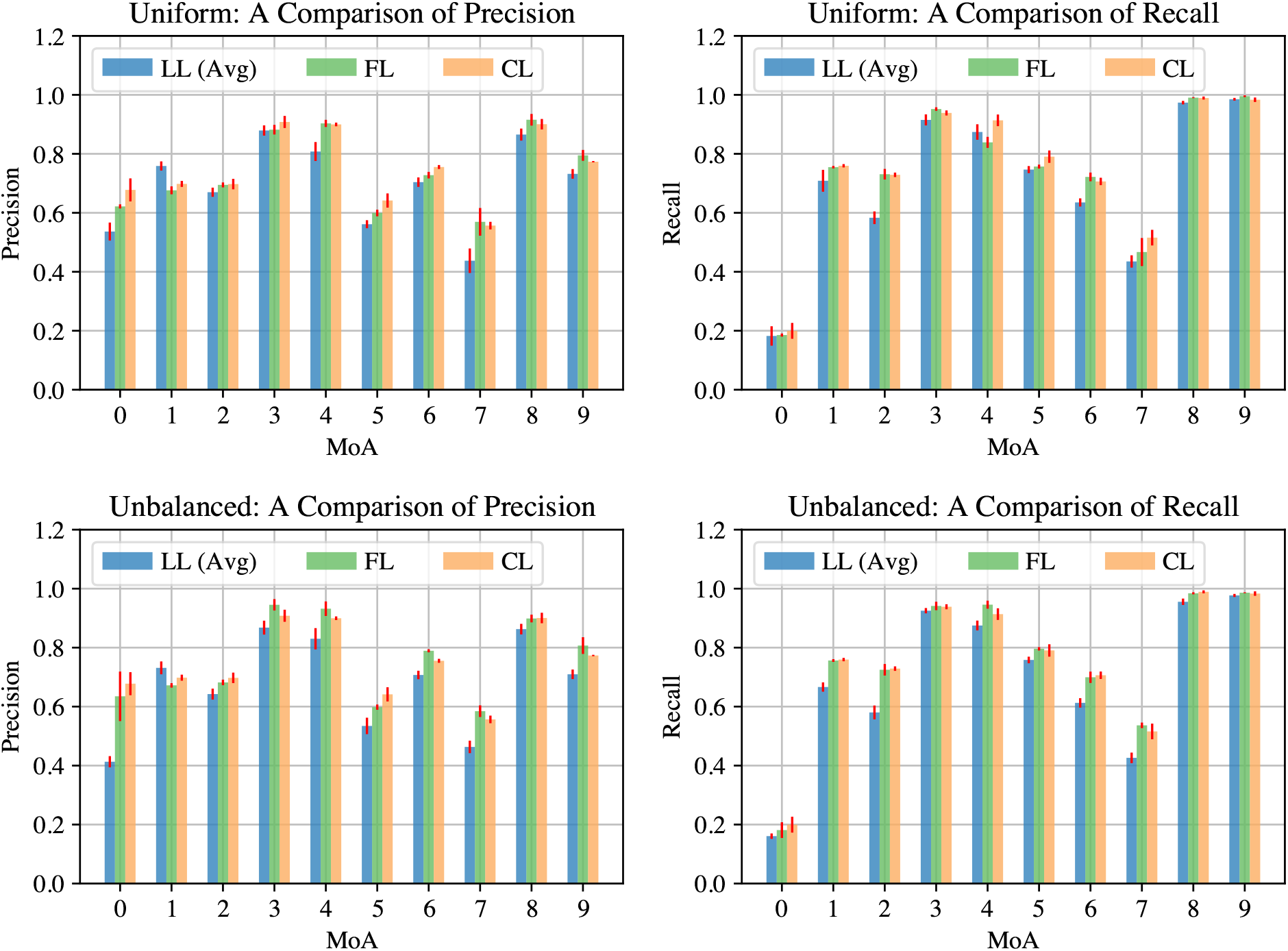
Comparison of Precision and Recall of Individual MoAs. This figure provides a comparison of precision and recall for each individual MoA among models trained under different schemes in both the uniform and unbalanced setups. Metrics for models trained with federated learning and centralized learning are represented as FL and CL, respectively. For a concise comparison, the metrics of local models are averaged and denoted as LL (Avg). **Left:** The two panels on the left compare precisions of different models in the uniform and unbalanced setups for each individual MoA. **Right:** The two panels on the right compare recalls of different models in the uniform and unbalanced setups for each individual MoA.

While inherent variations in precisions and recalls across different MoAs are expected, it’s noteworthy that these variations consistently follow similar trends across various models. Federated models consistently outperform local models in both precision and recall for the majority of individual MoAs, while exhibiting equivalent performance for a minority of MoAs. While the local model achieves exceptionally high precision for the protein synthesis inhibitor (encoded as MoA.1), it is important to note that this is accompanied by a lower recall, implying the potential bias of local training with insufficient data [56]. Furthermore, no significant statistical difference is observed in both precision and recall between federated and centralized models for any individual MoA.

We summarize that in both the uniform and unbalanced setups, federated learning demonstrates its ability to improve prediction performance across the majority of individual MoAs while showing minimal fairness concerns. Notably, when compared to centralized learning, federated learning achieves nearly identical prediction performance across individual MoAs. These results show the effectiveness of federated learning in improving predictive accuracy while preserving data privacy.

### 3.2 The More Participants, the Better Prediction Performance

Further we study the performance of federated models with respect to the number of participants. To minimize the influence of variation in statistical distributions of local datasets, the experiment is conducted on the uniform and unbalanced setup involving a total of 6 clients with a base model VGG13. With all hyperparameters for federated training controlled, we compare the performance of federated models trained with increasing numbers of participants. The experimental results are shown in Figure 6.

**Figure 6.**
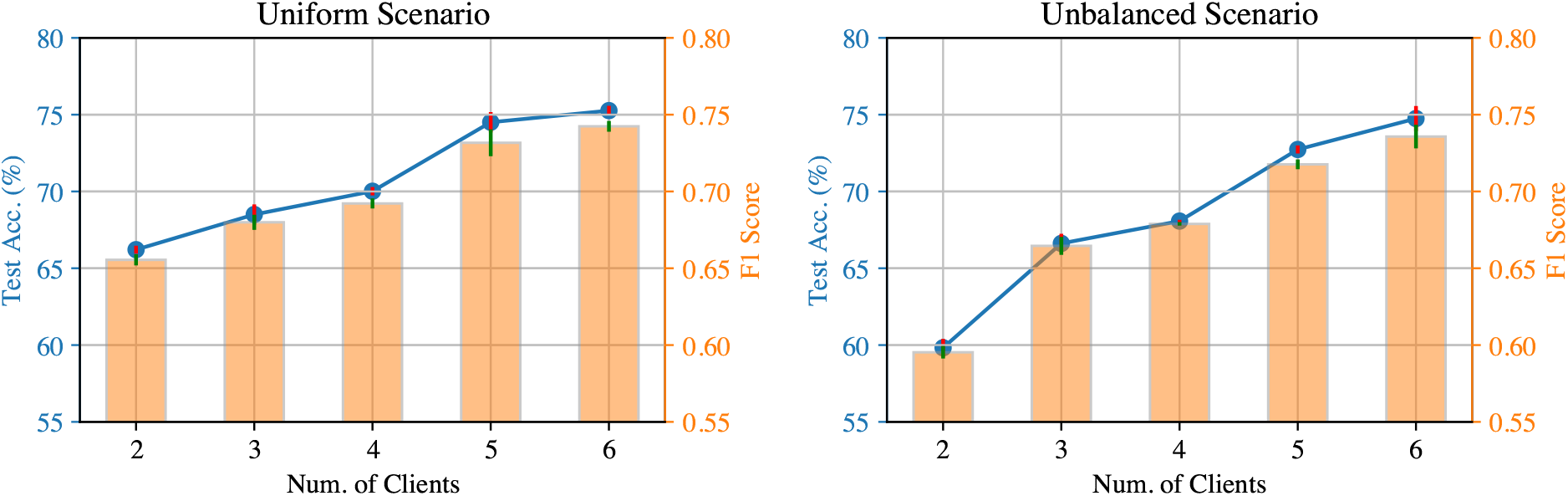
Performance of federated models trained with *k* ∈ {2, 3, 4, 5, 6} clients in the uniform and unbalanced scenarios. Test accuracy and macro F1 scores across MoAs are presented in blue curves and orange bars, respectively.

It is observed that with local datasets of approximately the same sizes in the uniform setup, with the number of participants increasing from 2 to 6, the test accuracy of federated models improves from 65.8% to 77.2%. The improvement of prediction ability is also reflected on the macro F1 scores of the models, which increases from 0.65 to 0.77. Consistent results are observed with the unbalanced setup, in which participants own local datasets of identical statistical distributions and different sizes. The involvement of more participants in the federated training improves the test accuracy and F1 scores of the federated model.

Therefore, we conclude that under the assumption that all participants possess data with homogeneous statistical distributions, a higher number of participants within the federated learning process leads to an improved prediction performance from the federated model.

### 3.3 Specialized Participant Brings Benefits

Within the pharmaceutical industry, a prevalent practice involves certain companies undertaking smaller-scale specialized studies on compounds associated with specific MoAs. The impacts of the involvement of these highly specialized clients in federated learning are yet to be studied, especially considering the possible deterioration of the model performance caused by data heterogeneity of local datasets. In this section, we compare the performance exhibited by federated models under two scenarios: one where the specialized client is excluded (FL-excluded), and the other with it included (FL-included). To quantify the effects, we evaluate both test accuracy for the specialized MoA and all MoAs of the models. The datasets and training scheme used in the experiment are shown as Figure 7. It can be seen that Client 0 is highly specialized on one specific MoA, while only a small fraction data The empirical results are presented in Figure 8. It is observed that when the public dataset comprises only 5% of MoA-related compounds, and the remainder is retained within the specialized participant’s local dataset, the federated model trained with FL-excluded manifests a test accuracy of 30.1% for MoA.s, while the FL-included model achieves a significantly higher average prediction accuracy of 68.3%. As the proportion of specialized data within the public dataset grows, both FL-excluded and FL-included models demonstrate improved test accuracies for the specialized MoA. However, despite this increase, the gap in performance persists. Even when 35% of compounds associated with the specialized MoA are included in the public data, the disparity between the two approaches remains statistically significant. While improving the prediction ability for the specialized MoA, models trained with FL-included encompass no significant deterioration of the prediction accuracy for other MoAs, which implies that the inclusion of specialised participants does not bring significant fairness problem for federated models.

**Figure 7.**
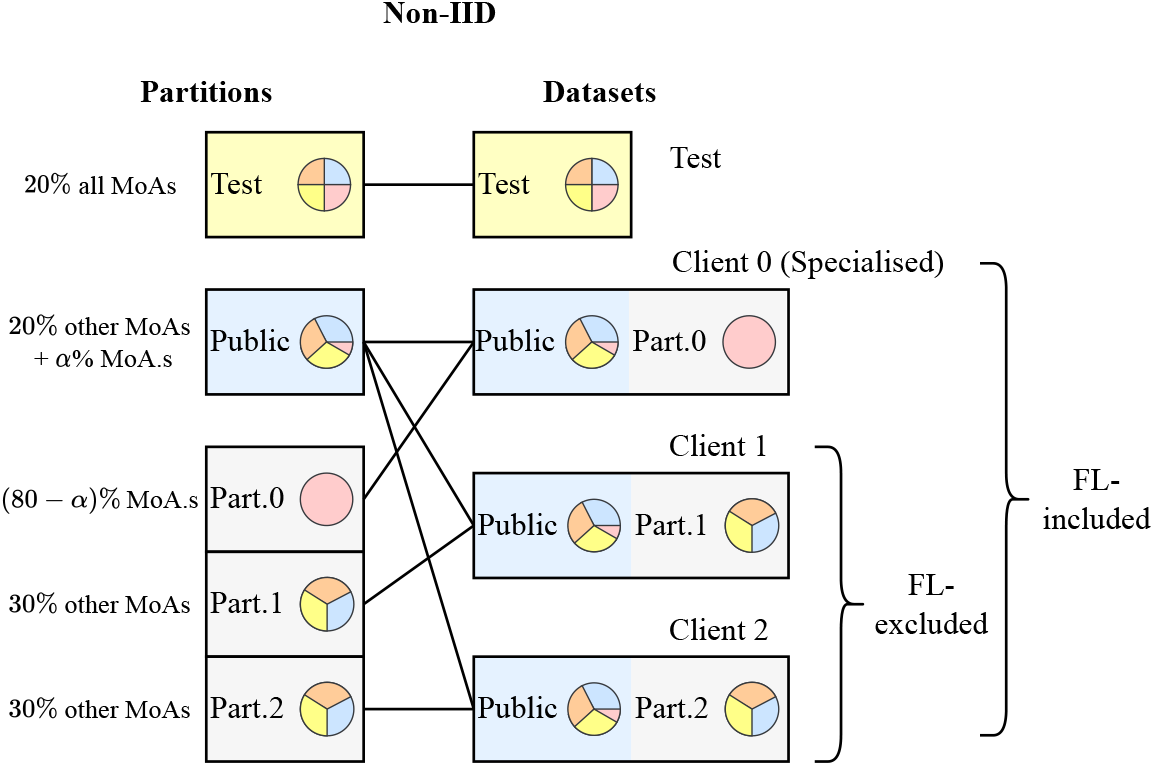
Partitioning and training scheme to simulate federated learning with specialized clients. Specifically, *α*% of MoA.s data is allocated to the public partition, while the remaining (80− *α*)% exclusively resides in a single private partition. Data of other MoAs are evenly distributed among the remaining private partitions. Two training schemes are compared with the specialised client included and excluded in the federated training, i.e. FL-included and FL-excluded.

**Figure 8.**
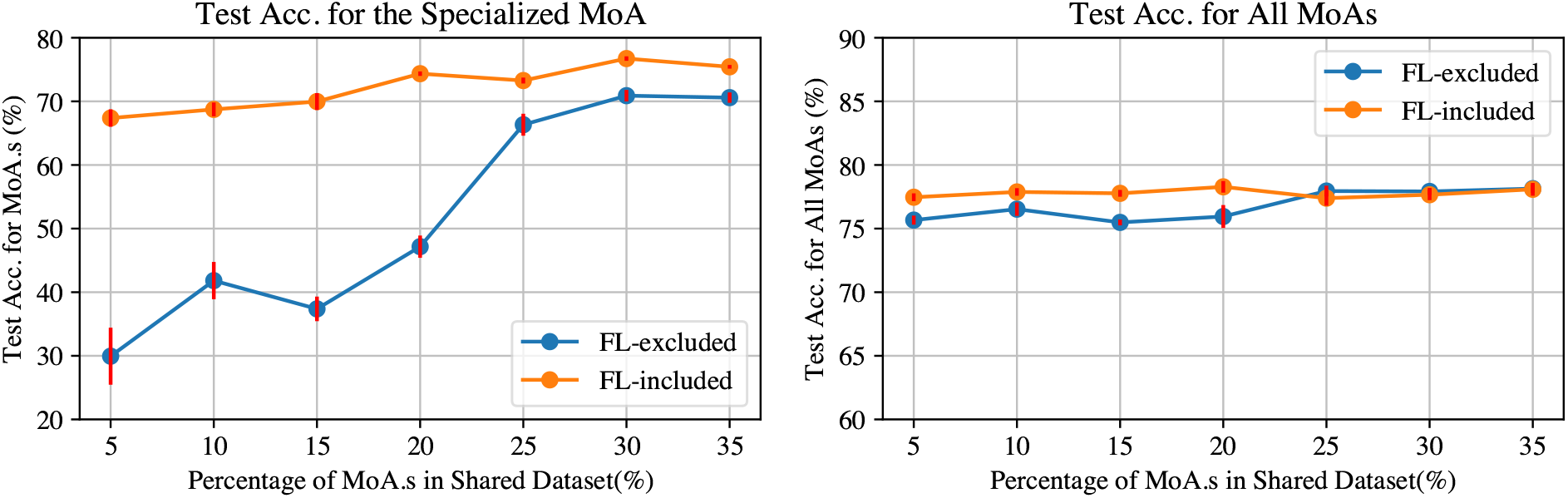
Comparison of Federated Model Performance in the Non-IID Setup. This figure compares the prediction accuracy of federated models in the non-IID setup when trained both with the specialized participant excluded (blue curves) and included (orange curves). **Left:** This panel displays the test accuracy for the specialized MoA (i.e. MoA.s) of federated models, comparing those trained with the specialized participant excluded and included. **Right:** This panel displays the test accuracy for all the MoAs of federated models, comparing those trained with the specialized participant excluded and included.

Thus, we summarize that the inclusion of the specialized client imparts a positive impact on the predictive capabilities for the specialised MoA and the fairness of the federated model. Moreover, as the availability of specialized MoA data within the public dataset diminishes, the specialized client’s contribution increasingly amplifies the model’s performance improvement.

## 4 Discussions and Conclusions

When assessing the mechanisms or safety of chemical compounds, traditional modeling based on chemical structure and machine learning (structure-activity relationships) are common. Prior research on federated learning in drug discovery and chemical safety assessment has focused on such chemical structure data [57, 58, 59]. However, chemical space is very large and using molecular or cellular data (such as high-content imaging experiments with Cell Paitning) has potential to better capture biological activity space, and hence improve predictions. A key challenge is however access to sufficient high quality data to be able to train accurate predictive models.

In this manuscript we applied federated learning to demonstrate that cross-organization collaborations are possible also for experiments with image-based readouts.

Our experiments offer empirical evidence that federated learning can provide several advantages for MoA predictions. We can summarize the findings based on the individual roles of different potential participants in the federation:

- Prospective participants: An organization not yet participating in a federation could, by joining an existing federated learning process, obtain a federated model with significantly improved performance compared to their local model, without the need to disclose their local data.
- Existing/founding participants: Existing participants within the federation benefits from remaining engaged in the federated learning process throughout the life cycle of a model. This is particularly important as new participants may continue to join over time, potentially leading to the development of federated models with even better future performance.
- Highly specialized participants: Participants with data related only to specific MoAs can also benefit from engaging in the federated learning process. Their involvement not only yields federated models with notably enhanced prediction performance for other non-specialized MoAs not available to them, but also contributes novel knowledge to the federation, significantly improving prediction accuracy for their specialized MoA.

From a broader perspective, by leveraging federated learning we show that it is possible to train models from isolated local data with equivalent performance to those trained with all data collected and centralized - something that is often infeasible in reality.

A common concern in federated learning is that statistical heterogeneity across local data could lead to performance issues, including convergence problems and fairness issues [60, 61, 62]. However, we argue that federated learning remains advantageous for MoA predictions for two key reasons. Firstly, the reported performance loss of federated models is compared against models trained with centralized data. However, federated learning models still outperform local models in principle, which is consistent with our experimental results. Secondly, the impact of statistical heterogeneity on federated learning may be limited. Recent arguments suggest that the key assumption of bounded gradient dissimilarity in previous theoretical analyses is too pessimistic to characterize data heterogeneity in practical applications and the performance loss is of little significance in many real-world federated training tasks [63] This is also consistent with our experimental results, where models trained with federated learning do not underperform centralized models with statistical significance.

In the context of drug discovery, by decoupling the joint distribution *p*_*i*_(***x***, *y*) = *p*_*i*_(***x***|*y*)*p*_*i*_(*y*), it becomes evident that data heterogeneity can be categorized into two main types:

- *p*_*i*_(*y*) (also known as label skew): Different participants may exhibit varying distributions of labels *y*. This variability can be attributed to different specializations among the participants.
- *p*_*i*_(***x***|*y*) (also known as feature skew): Different participants may display diverse distributions of data ***x***, even when the label *y* is fixed. This variability may arise due to variations in data collection protocols, system noises, or measurement methods among participants.

In our work, we have focused on the case of label skew, conducting empirical experiments. Meanwhile, there are other works that have explored feature skew in the context of drug discovery [58, 59]. However, the comprehensive impact of data heterogeneity in this domain remains an area for further investigation. Another limitation of our study is that we only include images generated from a single microscope. A more relevant scenario would be that images come from different microscopes but the same agreed-upon protocol (e.g. Cell Painting), and as more data is deposited in public databases this type of study will likely soon be possible to realize in future works. Another limitation is the number of tool compounds with high quality MoA annotations that are available to be tested experimentally, and this remains is a general challenge for the field. Efforts to develop such tool compounds are highly needed to further improve accuracy for prediction of MoAs.

In conclusion, we constructed hypothetical collaboration scenarios for organizations involved in assessing MoA for chemical compounds based on multiplexed high-content imaging experiments (Cell Painting) and demonstrated that federated learning enables training accurate models without disclosing data between collaborators. It is our hope that our findings will inspire to more multi-institutional collaborative machine learning initiatives.

## Acknowledgments

We acknowledge funding from the Centre for Interdisciplinary Mathematics at Uppsala University, the Swedish strategic initiative on e-Science eSSENCE, Swedish Research Council (grants 2020-03731 and 2020-01865), FORMAS (grant 2022-00940), Swedish Cancer Foundation (22 2412), and Horizon Europe grant agreement #101057014 (PARC) and #101057442 (REMEDI4ALL).

## Notes

### Competing Interest Statement

The authors have declared no competing interest.

